# Nanodomain formation in lipid bilayers II: The influence of mixed-chain saturated lipids

**DOI:** 10.1101/2025.10.28.685012

**Authors:** Deeksha Mehta, Emily H. Chaisson, Averi M. Cooper, Maryam Ahmed, M. Neal Waxham, Frederick A. Heberle

**Author notes:** These authors contributed equally (Deeksha Mehta, Emily H. Chaisson).

## Abstract

An important class of lipids found in biological membranes are composed of two structurally different hydrocarbon chains. Among these, low-melting lipids possessing both a saturated and unsaturated chain have been intensely studied because of their biological abundance and influence on lipid rafts. In contrast, much less is known about the biophysical effects of mixed chains in high-melting lipids. Here, we investigated two such lipids—MSPC (14:0-18:0 PC) and SMPC (18:0-14:0 PC)—to determine how chain length mismatch and acyl chain position on the glycerol backbone influence lateral organization. We studied the temperature-and composition-dependent phase behavior of liposomes composed of either mixed-chain or symmetric-chain high-melting lipids plus DOPC and cholesterol, using techniques sensitive to domain formation at both microscopic and nanoscopic length scales. All studied mixtures exhibited liquid-ordered (Lo) + liquid-disordered (Ld) phase coexistence with domains that were visible in confocal microscopy experiments. FRET measurements showed that all mixtures also exhibited nanoscopic heterogeneity at temperatures above the microscopic miscibility transition temperature, and cryo-EM imaging further revealed bilayer thickness variation consistent with coexisting Ld and Lo phases. Both the microscopic miscibility transition temperature, μm-*T_mix_*, and its nanoscopic counterpart, nm-*T_mix_*, were strongly correlated with the melting transition temperature of the saturated lipid; the sole exception was SMPC/DOPC/Chol, whose μm-*T_mix_* showed a significant negative deviation from the expected value, implying an enhanced propensity for nanoscopic phase separation in mixtures containing this high-melting species. These results point to strong effects of acyl chain position within mixed-chain high-*T_M_* lipids on the microscopic phase behavior of ternary mixtures.

## 1. Introduction

Lateral lipid heterogeneity is a fundamental feature of eukaryotic plasma membranes [1, 2], playing a role in cellular processes such as protein trafficking, signaling, and pathogen-host interactions [3, 4]. Although the existence of lipid rafts is strongly supported by abundant biochemical and spectroscopic evidence, direct visualization of raft domains has been challenging due to their small size and transient lifetimes [5], and studies using simplified model membrane mixtures have been crucial in establishing the physicochemical principles underlying raft formation [6, 7].

The most widely studied models for the plasma membrane outer leaflet are composed of cholesterol together with two phospholipids that have a large difference in their chain melting transition temperatures, *T_M_* [8]. The high-*T_M_* lipid is often a sphingomyelin or fully saturated PC lipid with 14 or more carbons in each chain, while the low-*T_M_* lipid typically has cis double bonds or methyl branches in one or both chains. When the low-*T_M_* lipid contains unsaturations in both chains (the typical example being DOPC), micron-sized domains are observed in giant unilamellar vesicles (GUVs) imaged with fluorescence microscopy [9]. However, lipids that have two unsaturated chains are relatively rare in biological membranes, with most naturally-occurring low-*T_M_* lipids instead having a saturated *sn*-1 chain (POPC being a typical example) [10]. Remarkably, substituting DOPC with POPC in a ternary mixture often results in GUVs that appear optically uniform [11, 12]. Nonetheless, nanoscopic lateral heterogeneity can be detected in these mixtures using techniques with appropriate spatial resolution such as Förster resonance energy transfer (FRET) [13, 14], electron spin resonance (ESR) [15, 16], small-angle neutron scattering (SANS) [17], and cryogenic electron microscopy (cryo-EM) [18, 19].

While studies such as those mentioned above have explored how mixed acyl chains in the low-*T_M_* lipid influence phase behavior, much less is known about the influence of mixed-chain high-*T_M_* lipids. Although the most abundant high-*T_M_* lipids in cell membranes typically have acyl chains of similar length [20], substantial differences in chain length are also observed. In some cases, this difference can be extreme; for example, sphingomyelin species with N-acyl chains as short as two carbons and as long as 28 carbons have been identified [21, 22]. In some yeast, saturated mixed-chain PC lipids are found in high concentrations and are upregulated in stressful conditions, suggesting a biological role [23]. Comparing saturated mixed-chain PC lipids with the same total number of acyl chain carbon atoms but different chain lengths—for example, MSPC (14,18-PC), DPPC (16,16-PC), and SMPC (18,14-PC)—the mismatched lipids have lower melting points than the symmetric species, with further differences seen when comparing two mismatched counterparts (i.e., SMPC and MSPC) [24]. Interestingly, single-component bilayers composed of MSPC or SMPC also show marked differences in interdigitation [25] and bending modulus [26], highlighting the importance of understanding how acyl chain positioning within mixed-chain lipids influences membrane biophysical properties.

Here, we focus on the behavior of MSPC and SMPC in ternary mixtures containing DOPC and cholesterol. To isolate the effects of chain mismatch and acyl chain positioning, we also examined mixtures containing symmetric-chain saturated lipids. We used fluorescence microscopy and cryo-EM to image vesicles with microscopic and nanoscopic spatial resolution, respectively. We further used fluorescence microscopy and FRET to measure the microscopic and nanoscopic miscibility transition temperatures (μm-*T_mix_* and nm-*T_mix_*, respectively). We find a strong correlation between these miscibility transition temperatures and *T_M_* of the saturated lipid except for SMPC/DOPC/Chol, whose μm-*T_mix_* occurs at a substantially lower temperature than expected.

## 2. Materials and Methods

### 2.1 Materials

Phospholipids 1,2-dioleoyl-sn-glycero-3-phosphocholine (DOPC), 1,2-dimyristoyl-*sn*-glycero-3-phosphocholine (DMPC), 1,2-dipentadecanoyl-*sn*-glycero-3-phosphocholine (15:0-PC), 1,2-dipalmitoyl-*sn*-glycero-3-phosphocholine (DPPC), 1,2-distearoyl-*sn*-glycero-3-phosphocholine (DSPC), 1-myristoyl-2-stearoyl-*sn*-glycero-3-phosphocholine (MSPC), 1-stearoyl-2-myristoyl-*sn*-glycero-3-phosphocholine (SMPC), and 1-palmitoyl-2-oleoyl-sn-glycero-3-[phospho-rac-(1-glycerol)] (sodium salt) (POPG) were purchased from Avanti Polar Lipids (Alabaster, AL, USA). Cholesterol was purchased from Nu-Chek Prep (Elysian, MN, USA). Fluorescent dyes 1,2-dioleoyl-*sn*-glycero-3-phosphoethanolamine-N-(lissamine rhodamine B sulfonyl) (ammonium salt) (LRPE), 1,2-distearoyl-*sn*-glycero-3-phosphoethanolamine-N-(7-nitro-2-1,3-benzoxadiazol-4-yl) (ammonium salt) (NBD-DSPE), and 1-palmitoyl-2-(dipyrrometheneboron difluoride)undecanoyl-*sn*-glycero-3-phosphocholine (TopFluor-PC, TFPC) were purchased from Avanti Polar Lipids. The fluorescent dye 1,1′-dioctadecyl-3,3,3′,3′-tetramethylindodicarbocyanine, 4-chlorobenzenesulfonate salt (DiD) was purchased from ThermoFisher Scientific (Waltham, MA, USA) and naphtho[2,3-a]pyrene (naphthopyrene, Nap) was purchased from TCI America (Portland, OR, USA). All lipids and dyes were suspended in HPLC-grade chloroform and stored at −20 °C until use. Concentrations of phospholipid stocks were determined using a colorimetric inorganic phosphate assay [27]; the concentration of the cholesterol stock was determined gravimetrically. Dye concentrations were determined from absorbance measurements using the following extinction coefficients: 95,000 M^-1^cm^-1^ (LRPE); 23,800 M^-1^cm^-1^ (Nap); 96,904 M^-1^cm^-1^ (TFPC); and 249,000 M^-1^cm^-1^ (DiD).

Ultrapure water was obtained from a Milli-Q IC 7000 purification system (Millipore Sigma, Burlington, MA, USA).

### 2.2 Confocal fluorescence microscopy

Giant unilamellar vesicles (GUVs) were prepared using a modified electroformation procedure [28, 29]. Briefly, a mixture of lipids and probes was prepared in the desired ratio by dispensing stock solutions into a 13 mm × 100 mm glass culture tube containing 100 µL of chloroform. The solution was then spread onto two indium tin oxide (ITO)-coated glass slides on a hot plate set at 55 °C (Delta Technologies, Stillwater, MN, USA) and placed under a vacuum for 2 h. A chamber was formed from the two slides using an O-ring spacer filled with 600 µL of 100 mM sucrose solution and then placed into an aluminum holder heated to 55 °C. GUVs were formed by applying a 2.0 V peak-to-peak, 10 Hz AC waveform to the chamber for 2 h while the holder was maintained at 55 °C, followed by cooling to 22 °C at a rate of 2.66 °C per hour. GUVs were harvested from the chamber and stored in a plastic microfuge tube before imaging (typically on the same day).

Prior to imaging, 50 μL of freshly harvested GUVs in 100 mM sucrose was suspended in 1.5 mL of 100 mM glucose in a glass culture tube. After approximately 30 min, a 5 μL aliquot was taken from the bottom of the tube and sandwiched between a glass slide and a cover slip for microscopy observations. Imaging was performed with a Nikon C2+ point scanning system attached to a Nikon Eclipse Ti2-E microscope (Nikon Instruments, Melville, NY, USA) equipped with a Plan Apo Lambda 60×/1.4 NA oil immersion objective. An objective cooling collar maintained a sample temperature of 22 °C (Bioptechs, Butler, PA, USA). NBD-DSPE and LRPE were excited with 488 nm and 561 nm laser lines, respectively, with quarter-wave plates (ThorLabs, Newton, NJ, USA) inserted into the excitation path to correct for polarization artifacts.

To determine the microscopic miscibility transition temperature, μm-*T_mix_*, of GUVs composed of ternary mixtures, the sample temperature was controlled using a VaHeat heating stage (Interherence, Erlangen, Germany), with the objective cooling collar used to access temperatures below 22 °C for some mixtures. As shown in Fig. S1, a representative field of view with at least 9 phase-separated vesicles was selected for each composition. The temperature was then incrementally increased by 1 °C and, after waiting 30 s for equilibration, the number of uniform and phase-separated vesicles was counted. This process was repeated until all vesicles appeared uniform. To determine μm-*T_mix_*, the fraction of phase-separated vesicles vs. temperature was fit to a sigmodal function,

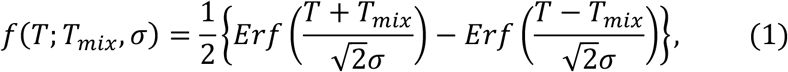

where 𝐸𝑟𝑓 is the general error function, 𝑇 is the temperature, and 𝑇*_mix_* and 𝜎 are parameters describing the center and width of the transition, respectively. 𝜎 and 𝑇*_mix_* were treated as free parameters to obtain best fit to the data. Error bars for 𝑇*_mix_* were calculated as the pooled standard error of the mean obtained from three sample replicates.

### 2.3 Fluorescence resonance energy transfer (FRET)

Vesicles of low average lamellarity (LLVs) for FRET measurements were prepared using the method of rapid solvent exchange (RSE) as previously described [30]. Stock solutions of lipids and probes in the desired ratios were dispensed into 13 mm × 100 mm glass screw-cap culture tubes containing 25 μL of chloroform. After adding 500 μL of ultrapure water, the tube was immediately mounted on the RSE device and exposed to vacuum while undergoing vigorous vortexing. During RSE, a slow leak of Ar was introduced into the tube to facilitate chloroform removal. After 1.5 min under vacuum, the sample was vented to argon and transferred to a plastic centrifuge tube.

FRET was measured using a HORIBA FluoroLog 3 spectrophotometer (HORIBA USA, Irvine, CA, USA) equipped with a temperature-controlled cuvette holder (Quantum Northwest, Liberty Lake, WA, USA). Samples contained three probes comprising two donor/acceptor FRET pairs, Nap/DiD and TFPC/DiD. Probe/lipid ratios were 1/200 for Nap, 1/1500 for TFPC, and 1/1000 for DiD. For fluorescence measurements, a 0.100 mL aliquot of a 0.625 mM (total lipid) sample was added to 1.90 mL of water in a fluorescence cuvette while stirring at 1000 rpm using a flea stir bar. Fluorescence intensity was measured in five channels (excitation/emission wavelengths in nm): Nap direct signal (427/545); Nap/DiD FRET (427/664); TFPC direct signal (499/551); TFPC/DiD FRET (499/664); and DiD direct signal (646/664). Excitation and emission slit widths were 2-3 nm, and the signal integration time was 0.1 s. Data were collected from 60 to 10 °C in 1°C decrements with an initial equilibration time of 15 min and a 2 min equilibration after each successive temperature change. The number of sample replicates was 10 for SMPC/DOPC/Chol, 13 for DPPC/DOPC/Chol, and 3 for MSPC/DOPC/Chol, 15:0-PC/DOPC/Chol, and DMPC/DOPC/Chol.

Sensitized acceptor emission was calculated from the intensity measured in the FRET channel after correcting for direct fluorescence contributions from donor and acceptor [16]. As described in the companion paper [31], the nanoscopic miscibility transition temperature, nm-*T_mix_*, was determined by fitting *FRET_R_* (calculated as the ratio of the FRET signals from the two probe pairs, i.e., TFPC/DiD FRET divided by Nap/TFPC FRET) to a phenomenological piecewise model consisting of a linear function at *T* > nm-*T_mix_* and a quadratic function at *T* < nm-*T_mix_*. Each dataset was also fit to a simpler quadratic function over the same temperature range to serve as a null model (i.e., the absence of a miscibility transition). The temperature range for data fitting was restricted to ∼ 10-15 °C on either side of the abrupt change in *FRET_R_* determined by visual inspection. For each fitted dataset, the piecewise function (i.e., corresponding to phase separation) was determined to be the most probable model as assessed by the difference in Akaike information content criterion as described in the companion paper [31].

### 2.4 Cryo-EM imaging

Large unilamellar vesicles (LUVs) were prepared as previously described [32]. Briefly, aqueous lipid dispersions with a total lipid concentration of 3 mg/mL were made by first mixing the required volumes of lipid stocks in chloroform using a glass Hamilton syringe (Hamilton Co., Reno, NV, USA). The solvent was then evaporated using an inert gas stream, and the sample was kept under vacuum overnight. Dried lipid films were hydrated with ultrapure water preheated to 55 °C and incubated for ∼1 h with vortex mixing every 15 min, followed by five freeze/thaw cycles between liquid nitrogen and a 55 °C water bath. This suspension was then extruded 31 times through a 100 nm polycarbonate filter using a handheld mini-extruder (Avanti Polar Lipids) maintained at 55 °C. The size and polydispersity of the LUVs were determined using dynamic light scattering (LiteSizer 100, Anton Paar U.S.A., Vernon Hills, IL, USA) immediately after preparation and again before cryo-preservation, which was typically performed 1 day after LUV preparation.

Cryo-preservation was performed by adding 4 μL of LUVs to a Quantifoil 2/2 carbon-coated 200 mesh copper grid (Electron Microscopy Sciences, Hatfield, PA, USA) that was glow-discharged for 30 s at 20 mA in a Pelco Easi-Glow discharge device (Ted Pella, Inc., Redding, CA, USA). This was followed by manual blotting at room temperature, in which the grids were plunged into liquid ethane cooled with liquid nitrogen. The cryo-preserved grids were subsequently stored in liquid nitrogen.

Cryo-EM image collection was performed at approximately 2 μm underfocus on a Titan Krios operated at 300 keV equipped with a Gatan K2 Summit direct electron detector operated in counting mode. Data collection was conducted in a semi-automated fashion using Serial EM software operated in low-dose mode. Briefly, areas of interest were identified visually, and 8×8 montages were collected at low magnification (2400×) at various positions across the grid, with desired areas marked for automated data collection. Data was collected at 2.7 Å/pixel. Movies of 30 dose-fractionated frames were collected at each target site with the total electron dose being kept to < 20 e^-^/Å^2^ to avoid beam-induced damage [33]. Dose-fractionated movies were drift-corrected with MotionCor2. Defocus and astigmatism were assessed with CTFFind4 [34]. Finally, a high-pass filter was applied along with phase flipping using the ‘mtffilter’ and ‘ctfphaseflip’ routines in the IMOD v4.11 software package.

Projection images of vesicles were analyzed in Wolfram Mathematica v. 13 (Wolfram Research Inc., Champaign, IL, USA) as previously described to obtain spatially resolved intensity profiles (IPs) in the direction normal to the bilayer [32]. Briefly, vesicle contours (i.e., the set of points corresponding to the midplane of the projected bilayer as defined by a relatively bright central peak) were first generated using a neural network-based algorithm (MEMNET) that is part of the TARDIS software package [35]. Vesicles meeting any of the following criteria were omitted from analysis due to the possibility of artifacts: (i) contact with the edge of the well; (ii) location at the edge of the image; (iii) containing internal debris, nested vesicles, or multiple lamellae; (iv) sufficiently non-spherical in shape; examples of selected and omitted vesicles can be seen in Fig. S2. For each selected vesicle, the MEMNET contour was resampled at arc length intervals of 5 nm, resulting in a polygonal representation of the 2D contour. For each polygon, all pixels within a 5 nm × 20 nm rectangular region of interest centered at the face were selected, and their intensities binned at 1 Å intervals in the long dimension (i.e., normal to the face) and subsequently averaged in the short dimension to produce a local segment IP. To determine the phase state of each bilayer segment, we subjected the local IP vectors to 2-means clustering using Mathematica’s FindClusters algorithm with the distance function set to Euclidean and all other parameters set to their default values. We also calculated the local bilayer thickness, *D_TT_*, as the distance between the two minima of the local IP. Two methods were used to locate the minima: (1) a ‘model-free’ method, in which a local 5-point Gaussian smoothing was first performed, and the distance between the two absolute minimum intensity values on either side of the central peak was determined; (2) a ‘model-fit’ method that fits the profile as a sum of four Gaussians and a quadratic background, with the troughs corresponding to the two absolute minimum intensity values on either side of the central peak. The two methods typically agree to within 1 Å; the average of the two measurements was taken as the raw segment thickness. The final reported segment *D_TT_* values were obtained by local 4-point Gaussian smoothing of the raw segment thickness values.

### 2.5 Differential scanning calorimetry (DSC)

Multilamellar vesicle samples for DSC were prepared by adding 1.0 mL of ultrapure water to ∼5 mg of dry lipid powder and then vortexing vigorously to disperse the lipid. The sample was hydrated at 55 °C for 2 h with intermittent vortexing, followed by five freeze/thaw cycles between −80 and 55 °C. DSC measurements were taken on a TA Nano DSC (TA Instruments, New Castle, DE, USA). The sample was first annealed with two complete temperature cycles between 5 and 60 °C at a scan rate of 1 °C/min. A production cycle was then collected from 5 to 60 °C at a scan rate of 0.1 °C/min.

## 3. Results and Discussion

Our primary goal was to determine how high-melting saturated lipids with mixed acyl chains (i.e., having different length) influence the phase behavior of ternary mixtures that are frequently used as models for the composition of the eukaryotic plasma membrane outer leaflet. We examined two such mixed chain lipids, MSPC and SMPC. Each of these lipids possesses fully saturated 14-carbon and 18-carbon chains, but the position of the chains on the glycerol backbone is reversed as shown in Fig. 1: the shorter chain is in the *sn*-1 position for MSPC and in the *sn*-2 position for SMPC. Because mixed-chain phosphatidylcholines can undergo limited acyl chain migration during synthesis, we independently verified the thermotropic behavior of MSPC and SMPC by differential scanning calorimetry (Fig. S3). The measured transition temperatures and thermogram shapes closely match those reported by Chen and Sturtevant [36], indicating a likely comparable (∼10%) level of positional isomerization and implying that the differences between MSPC and SMPC reported here are conservative estimates.

**Figure 1.**
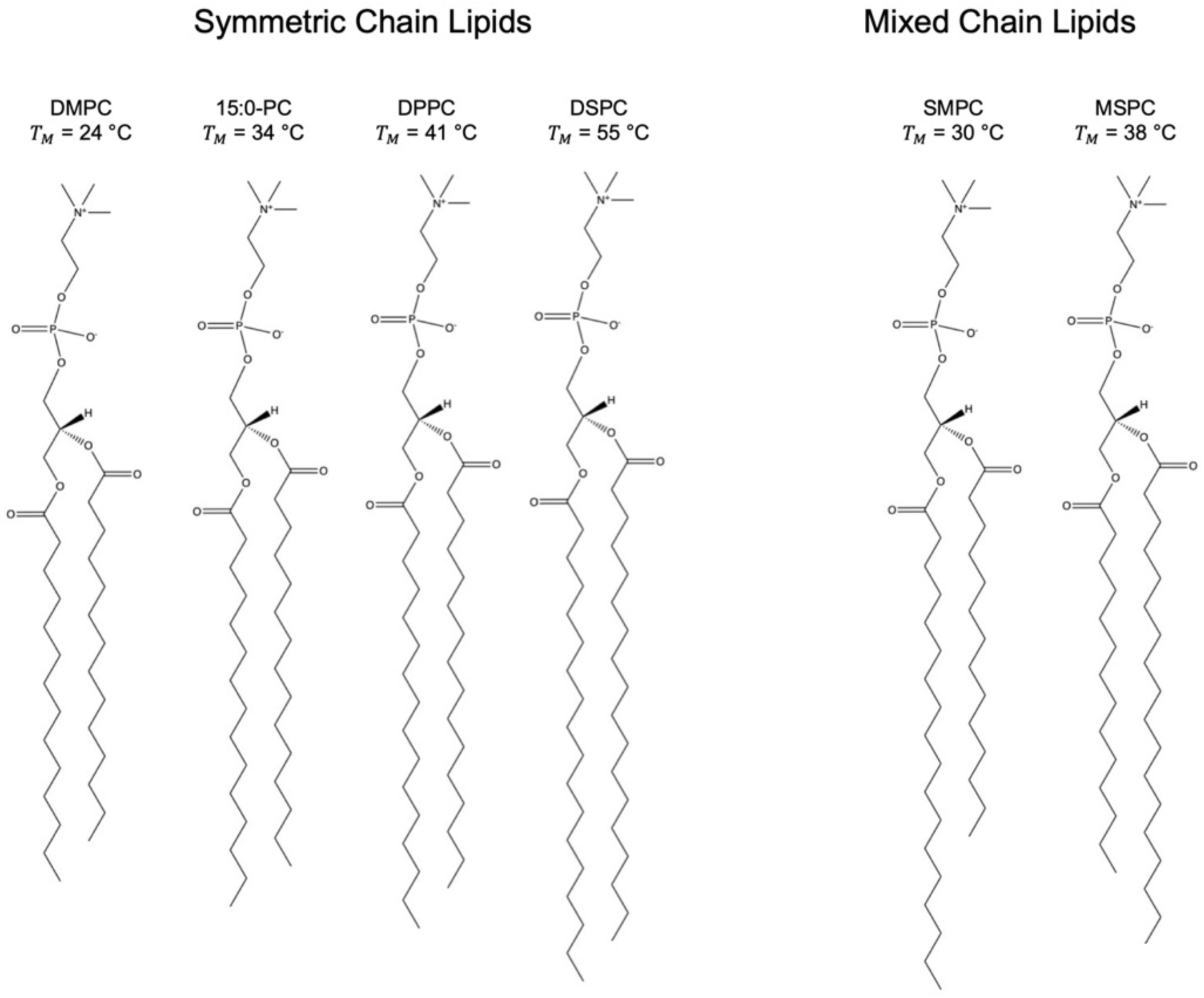
High-*T_M_* lipids in this study. From left to right: DMPC (14:0-14:0 PC); 15:0-PC (15:0-15:0 PC); DPPC (16:0-16:0 PC); DSPC (18:0-18:0 PC); SMPC (18:0-14:0 PC); and MSPC (14:0-18:0 PC). *T_M_* values are taken from Koynova and Caffrey [42].

The natural tilt of the glycerol backbone results in a vertical offset of approximately 0.5 Å between the *sn*-1 and *sn*-2 chains of fluid-phase PC bilayers, with the *sn*-2 chain residing at the higher position on average [25]; this backbone tilt slightly lessens the effective chain length mismatch of MSPC and slightly enhances it for SMPC. In addition to the high-*T_M_* lipid, each of the studied mixtures included cholesterol and the low-melting lipid DOPC. As a baseline for comparison, we also examined mixtures that contained 15:0-PC as the high-*T_M_* species, as this lipid has two identical chains and a main chain transition temperature between that of MSPC and SMPC [37].

At the molecular level, differences in acyl chain length and position are known to influence lipid packing, chain interdigitation, and bilayer thickness, particularly near the chain melting transition [36, 38, 39]. These microscopic features are expected to affect line tension and domain stability, and thus indirectly influence miscibility behavior. In the present study, however, we focus on experimentally observable miscibility transitions rather than attempting to resolve the underlying molecular packing or lipid dynamics directly. Our results therefore provide a window into the influence of chain asymmetry on lateral organization, complementary to prior structural and spectroscopic studies of mixed-chain phospholipids.

### 3.1 Microscopic phase behavior from fluorescence imaging

We used confocal fluorescence microscopy (CFM) to examine the microscopic phase behavior of GUVs composed of ternary mixtures of cholesterol, the low-*T_M_* lipid DOPC, and a high-*T_M_* lipid: MSPC, SMPC, or 15:0-PC. Figure 2 shows confocal slices near the GUV cap for representative GUVs composed of a 1:1 ratio of MSPC:DOPC (top row), 15:0-PC:DOPC (middle row), and SMPC:DOPC (bottom row) with varying cholesterol concentration as indicated at 22 °C. In the mixtures containing MSPC (top row) or 15:0-PC (middle row), the irregularly shaped, dark domains with rigid boundaries at low cholesterol concentrations (≤ 10 mol%) were suggestive of coexisting fluid and gel phases (i.e., Ld+Lβ). At 20 mol% cholesterol, the domain boundaries were smoother and showed an increased tendency for rounded shapes, behaviors that are consistent with coexisting liquid phases (i.e., Ld+Lo). At 40 mol% cholesterol, uniform mixing was observed for both compositions. In contrast to the behavior for MSPC and 15:0-PC, mixtures containing SMPC (Fig. 2, bottom row) showed no evidence of phase separation except at the lowest cholesterol concentration of 5 mol%, where GUVs had a distinctive patchy appearance.

**Figure 2.**
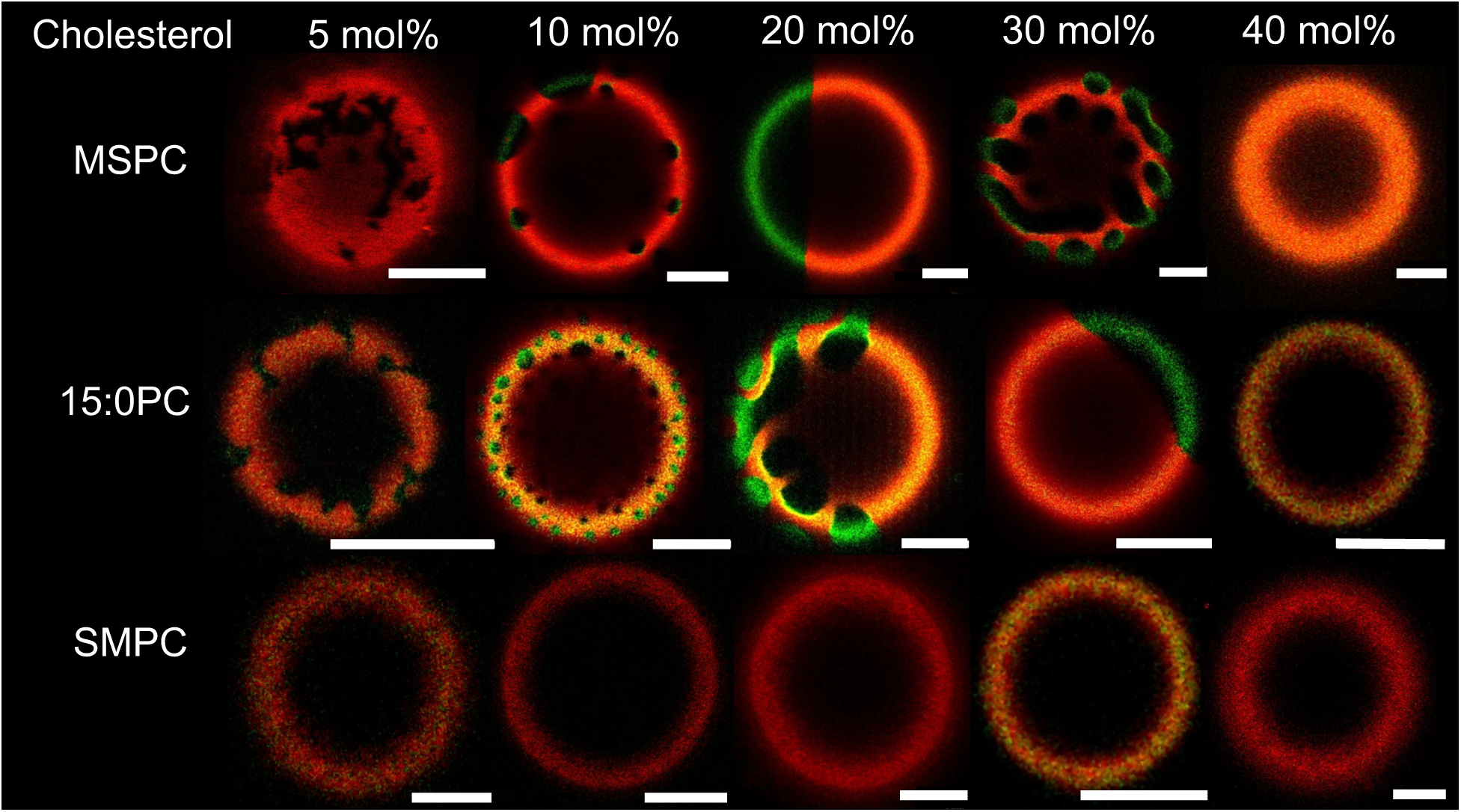
Confocal fluorescence images of GUVs at 22 °C. Shown are slices near the GUV cap for three-component mixtures containing 1:1 high-*T_M_*/DOPC with increasing Chol mol% and different high-*T_M_* lipid as indicated: MSPC (top row); 15:0-PC (middle row); and SMPC (bottom row). GUVs were labeled with fluorescent dyes LRPE (red) and NBD-DSPE (green). Each image shows the composite of the red and green channels. Scale bars are 5 µm.

Figure 3 summarizes the microscopic phase behavior of MSPC/DOPC/Chol, 15:0-PC/DOPC/Chol, and SMPC/DOPC/Chol as assessed from GUVs. The diagrams for MSPC/DOPC/Chol and 15:0-PC/DOPC/Chol (Fig. 3a and 3b, respectively) are similar and broadly consistent with other mixtures of saturated lipids, DOPC, and cholesterol [8], revealing regions of coexisting Ld+Lβ (blue triangles) and Ld+Lo (green circles) at 22 °C. Although we did not observe conclusive visual evidence of Ld+Lβ+Lo coexistence, we can infer from the Gibbs phase rule that a three-phase coexistence region must be present between the Ld+Lβ and Ld+Lo regions due to the absence of an intervening single-phase region. Interestingly, a qualitatively different diagram was found for SMPC/DOPC/Chol (Fig. 3c), which shows only a narrow region of Ld+Lβ below 5 mol% cholesterol and no evidence of Ld+Lo coexistence. These results point to a strong influence of acyl chain positioning within mixed chain high-*T_M_* lipids on the microscopic phase behavior of ternary mixtures.

**Figure 3.**
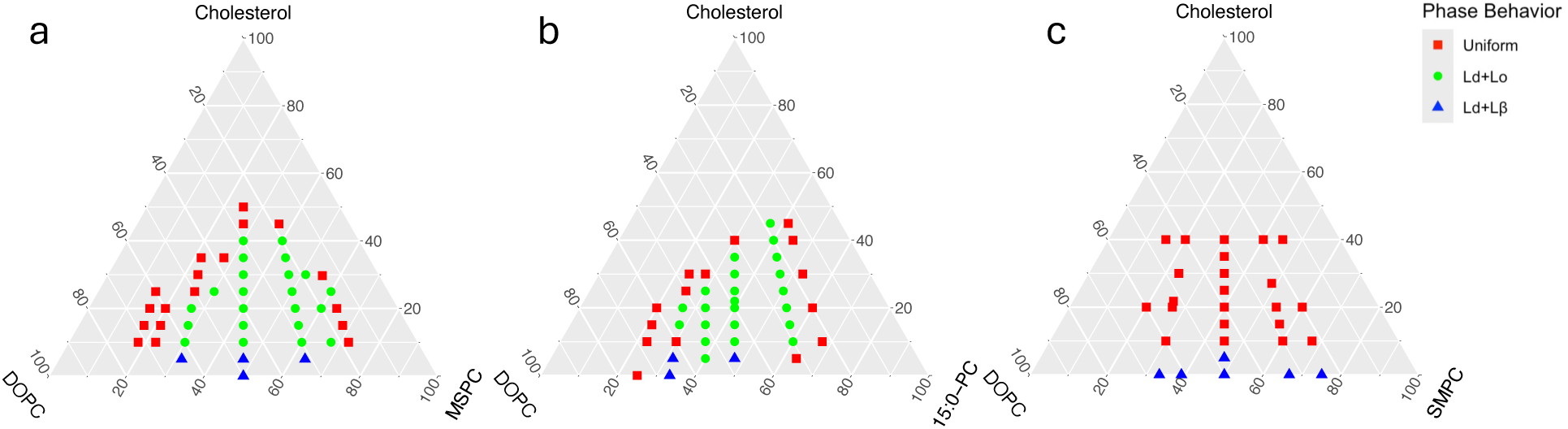
Phase behavior of GUVs at 22 °C. Shown are ternary phase diagrams for: MSPC/DOPC/Chol (a); 15:0-PC/DOPC/Chol (b); and SMPC/DOPC/Chol (c). Each point in the diagram shows the predominant phase behavior at a specific composition: Ld+Lβ (blue triangles), Ld+Lo (green circles), uniform mixing (red squares).

### 3.2 Microscopic miscibility transition temperature (μm-T_mix_) from fluorescence imaging

Although differential scanning calorimetry has been used to probe Ld+Lo miscibility in multicomponent lipid membranes [40], such transitions are typically associated with weak enthalpy changes and broad calorimetric features [41], making precise determination of a miscibility transition temperature challenging. As a result, microscopic miscibility transitions are more reliably quantified using optical methods that directly resolve domain formation. Accordingly, we determined μm-*T_mix_* of various ternary mixtures with the composition 40/40/20 mol% high-*T_M_* lipid/DOPC/Chol using confocal microscopy. After initially cooling the sample to 17 °C (where all systems showed coexisting Ld and Lo phases), temperature was increased until all vesicles were visibly uniformly mixed. Representative data for the fraction of phase separated vesicles vs. temperature is shown in Fig. 4a. We determined μm-*T_mix_* as the midpoint of a sigmoidal function fitted to each dataset, shown as solid lines in Fig. 4a. Except for SMPC/DOPC/Chol, the trend for μm-*T_mix_* follows that of the main chain melting transition temperature, *T_M_*, of the saturated lipid (Fig. 4b) [42].

**Figure 4.**
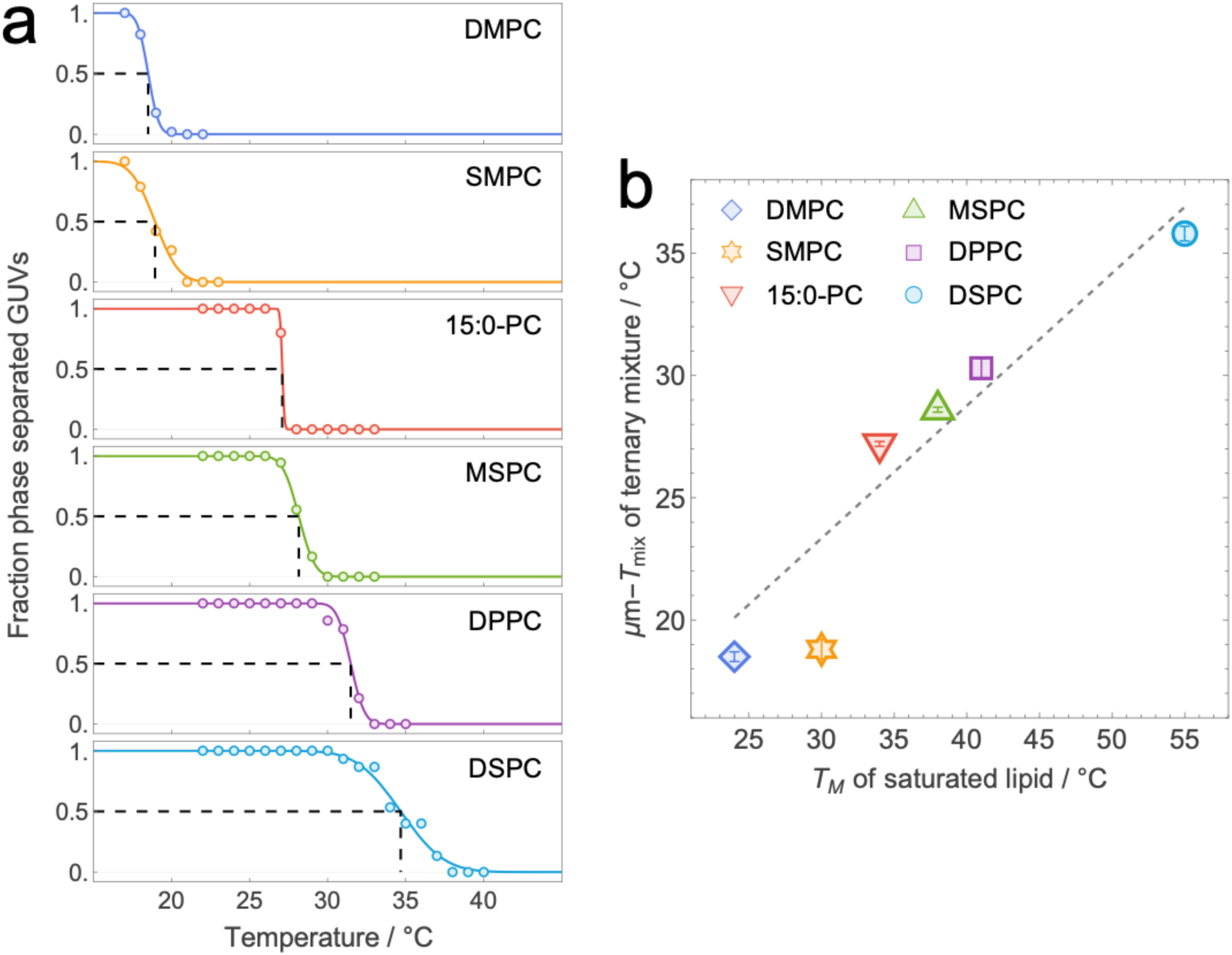
Microscopic miscibility transition temperature (μm-*T_mix_*) determined from confocal microscopy of GUVs. (a) The fraction of phase separated vesicles vs. temperature (open symbols) for ternary mixtures composed of high-*T_M_* lipid/DOPC/Chol 40/40/20 mol% for the indicated high-*T_M_* lipids. Data for one of three experimental replicates is shown overlaid with a fitted sigmoidal function (solid lines). The vertical dashed lines indicate the midpoint of the sigmoidal function, μm-*T_mix_*. (b) μm-*T_mix_* for ternary mixtures vs. the chain melting temperature, *T_M_*, of the saturated lipid. Average values and uncertainties obtained from sample replicates (*N* = 3) are listed in Table 2. The dashed line is a linear fit (R^2^ = 0.998) to the data points corresponding to symmetric chain high-*T_M_* lipids (i.e., DMPC, 15:0-PC, DPPC, and DSPC). Error bars are SEM determined from sample replicates. *T_M_* values are taken from Koynova and Caffrey [42].

### 3.3 Nanoscopic phase behavior from cryo-EM imaging

It is frequently observed that ternary mixtures containing mixed chain *low-melting* lipids exhibit nanoscopic lateral heterogeneity at physiologically relevant cholesterol concentrations [16]. In those cases, the low-*T_M_* lipid possesses a fully saturated *sn*-1 chain and one or more cis double bonds in the *sn*-2 chain. Although domains in such mixtures are smaller than the resolution limit of CFM (∼ 200 nm), their presence has been inferred from spectroscopic and scattering experiments. In principle, nanoscopic domains can also be directly visualized using cryo-EM [18, 43]. Unlike fluorescence microscopy, which requires partitioning of a probe molecule between the coexisting phases to distinguish them, the analogous contrast in cryo-EM images arises from intrinsic differences in the bilayer thickness and electron density of ordered and disordered phases [44].

Representative cryo-EM images of ternary mixtures (high-*T_M_* lipid/DOPC/Chol 40/40/20 mol%) are shown in Fig. 5, and full field of view images are shown in Fig. S2. Consistent with the microscopic Ld+Lo coexistence observed in GUVs, LUVs in which the high-*T_M_* lipid was DPPC, MSPC, or 15:0-PC (Fig. 5a, 5b, and 5c, respectively) showed coexisting regions with average thicknesses of 30-31 Å and 35-37 Å. These values are similar to the thicknesses of Ld and Lo phases measured with X-ray and neutron scattering [17, 45]. Remarkably, similar thickness variability was also observed for LUVs that contained SMPC (Fig. 5c) despite the uniform appearance of GUVs with this composition (Fig. 2). Schick and others have argued that nanodomains are properly described as a 2D microemulsion that originates in coupled fluctuations of local bilayer curvature and composition [46, 47]. The microemulsion theory predicts the existence of spatially modulated Ld-like and Lo-like domains with a characteristic length scale of ∼ 100 nm in mixtures where the saturated and unsaturated lipids have very different spontaneous curvatures. The theory further predicts that these domains are anti-registered (i.e., an Lo domain in one leaflet is always opposite an Ld domain in the other leaflet), which would presumably result in a nearly uniform overall membrane thickness in compositionally symmetric bilayers [48, 49]. Our cryo-EM images are inconsistent with this prediction, instead revealing substantial thickness variation within individual vesicles that suggests fully registered Ld and Lo phases; this is observed for each of the four compositions, including the SMPC/DOPC/Chol mixture that is visually uniform at the microscopic length scale at 22 °C (the cryo-preservation temperature).

**Figure 5.**
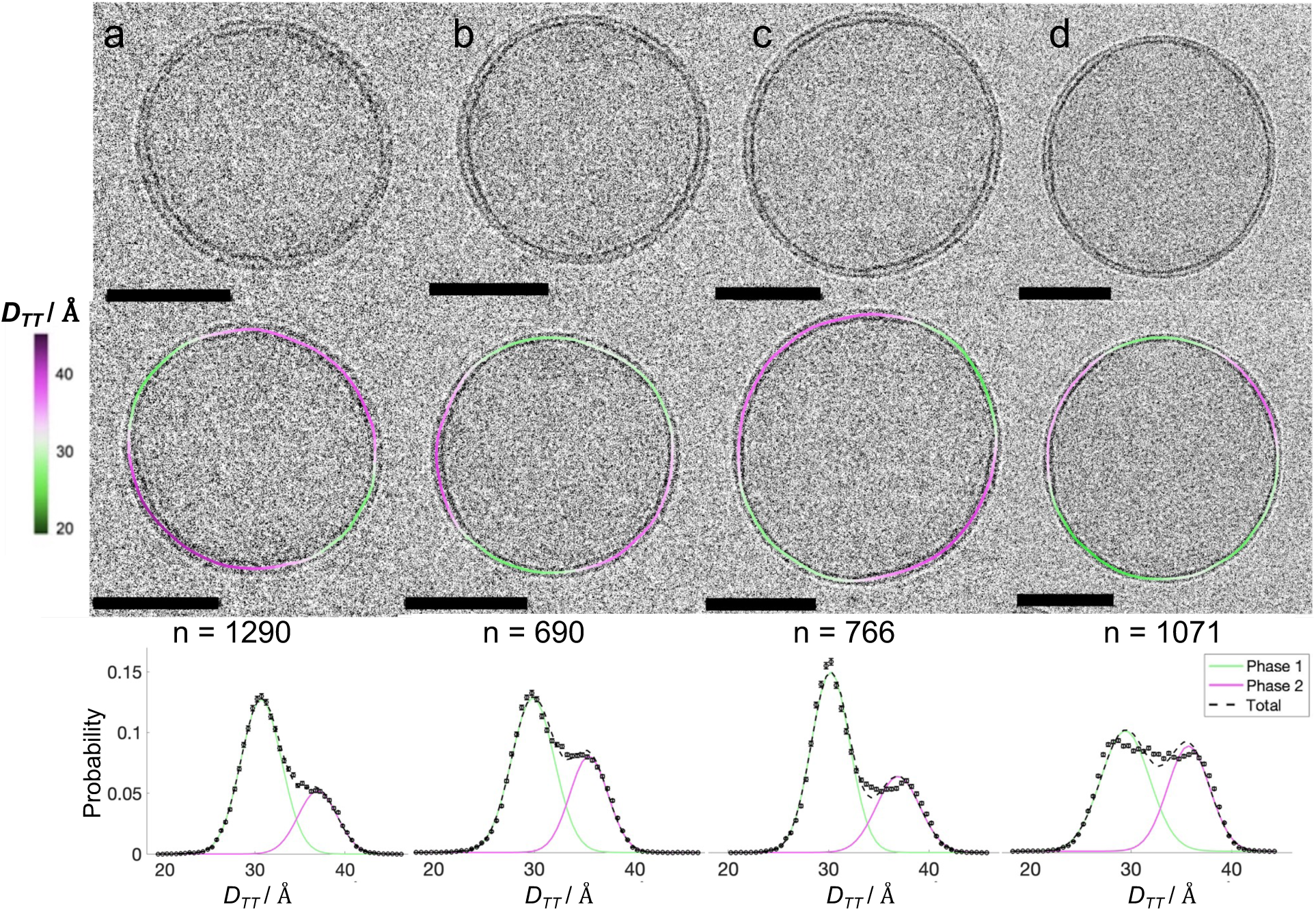
Cryo-EM images of ternary LUVs. All compositions are 40/40/20 mol%: DPPC/DOPC/Chol (a); MSPC/DOPC/Chol (b); 15:0-PC/DOPC/Chol (c); and SMPC/DOPC/Chol (d). Shown is a representative vesicle for each composition, without (top row) and with (middle row) an overlay of the local bilayer thickness, *D_TT_*, as indicated by the color scale bar. Black image scale bars are 50 nm. Plotted in the bottom row are histograms of local bilayer thickness calculated as described in Methods, with the total number of vesicles contributing to each histogram denoted by *n*.

Further insight can be gained from histograms of the local bilayer thickness obtained from the full population of vesicles (Fig. 5, bottom row). For each of the four mixtures, these histograms show a bimodal thickness distribution consistent with the thickness variability that is visually apparent in the image. Fitting the distributions as a sum of two gaussians yields values for the mean thickness and area fraction of each phase (Table 1). Notably, for all mixtures except SMPC/DOPC/Chol, the probability density peak is larger for the thinner Ld phase compared to the thicker Lo phase. In contrast, SMPC/DOPC/Chol has a smaller Ld peak and a greater probability of thicknesses between that of pure Ld or Lo. Intermediate thickness values likely correspond to domain boundaries where density from both phases contributes to the projected image. Because the frequency of boundary segments should increase with increasing domain perimeter in the vesicle, we speculate that SMPC promotes smaller domains with a concomitant increase in domain interface.

**Table 1.** Analysis of LUVs from cryo-EM images.

| Mixture<br>(40/40/20 mol%) | $D_{TT} / \text{\AA}$ | | Lo area fraction |
| --- | --- | --- | --- |
|  | Lo | Ld |  |
| DPPC/DOPC/Chol | $37.0 \pm 2.2$ | $30.7 \pm 2.3$ | 0.28 |
| MSPC/DOPC/Chol | $36.9 \pm 2.2$ | $30.2 \pm 1.9$ | 0.36 |
| 15:0-PC/DOPC/Chol | $35.4 \pm 1.9$ | $29.9 \pm 2.6$ | 0.36 |
| SMPC/DOPC/Chol | $35.7 \pm 2.1$ | $29.4 \pm 2.4$ | 0.43 |

**Table 2.** Miscibility transition temperatures for ternary mixtures (error is SEM).

| Mixture (40/40/20 mol%) | $\mu\text{m-}T_{mix} / ^\circ\text{C}$ | $\text{nm-}T_{mix} / ^\circ\text{C}$ |
| --- | --- | --- |
| DMPC/DOPC/Chol | $18.5 \pm 0.2$ | $21.44 \pm 0.03$ |
| SMPC/DOPC/Chol | $18.8 \pm 0.4$ | $28.6 \pm 0.2$ |
| 15:0-PC/DOPC/Chol | $27.2 \pm 0.1$ | $30.3 \pm 0.4$ |
| MSPC/DOPC/Chol | $28.6 \pm 0.1$ | $31.6 \pm 0.1$ |
| DPPC/DOPC/Chol | $30.3 \pm 0.4$ | $35.4 \pm 0.1$ |
| DSPC/DOPC/Chol | $35.8 \pm 0.3$ | $45.1 \pm 0.5$ |

### 3.4 Nanoscopic miscibility transition (nm-T_mix_) temperature from FRET spectroscopy

To more fully characterize the phase behavior of the 40/40/20 mol% samples, we used FRET to determine nm-*T_mix_*. As described in the companion paper [31], FRET is sensitive to changes in the spatial organization of donor and acceptor lipid probes that occur when the bilayer undergoes a miscibility transition. Provided these probes partition non-uniformly between the coexisting phases, the FRET signal changes abruptly upon formation of domains whose size exceeds approximately twice the Förster distance, *R_0_*, of the donor-acceptor pair. As *R_0_* for lipid fluorophores is typically in the range of 2-6 nm, FRET can detect the formation of domains as small as 5-10 nm that cannot be detected by conventional optical microscopy.

Our experiments utilized three fluorescent lipid probes comprising two donor-acceptor pairs: (1) Nap donor to DiD acceptor; and (2) TFPC donor to DiD acceptor. Co-localization of TFPC and DiD within the Ld phase results in enhanced FRET efficiency at temperatures below nm-*T_mix_*. However, because Nap preferentially partitions to the Lo phase, FRET between Nap and DiD decreases upon Ld+Lo phase separation (Fig. S4). The use of two probe pairs with complementary partitioning behaviors in the same sample allows us to distinguish between changes in probe spatial organization due to phase separation and other, confounding factors that can also influence FRET efficiency, such as temperature-dependent variation in molecular packing density that would be expected to change the two FRET signals in a similar way. For quantitative analysis, we fit the ratio of the two FRET signals (i.e., TFPC:DiD / Nap:DiD) to a phenomenological piecewise model that captures the different behavior of the regimes above and below nm-*T_mix_* as described in the companion paper [31]. We find that analyzing the ratio of the FRET signals provides a more robust determination of nm-*T_mix_*.

The FRET ratio as a function of temperature, plotted in Fig. 6a, shows a complicated behavior as the temperature is lowered from 60 °C that suggests the presence of a miscibility transition. For each mixture, the signal is relatively flat at higher temperature but increases abruptly upon the formation of Lo domains at the nanoscopic miscibility transition temperature. The solid lines in Fig. 6 show the best fit curve corresponding to a piecewise model of uniform mixing in the high temperature regime and phase separation in the low temperature regime, with the short vertical bar indicating the temperature that separates these regimes (i.e., nm-*T_mix_*). For the mixtures with the highest values of nm-*T_mix_*, the signal plateaus and eventually decreases as temperature is quenched deeper into the Ld+Lo region. As shown in Fig. 6b and Table 2, nm-*T_mix_* of the ternary mixtures increases linearly with *T_M_* of the saturated lipid.

**Figure 6.**
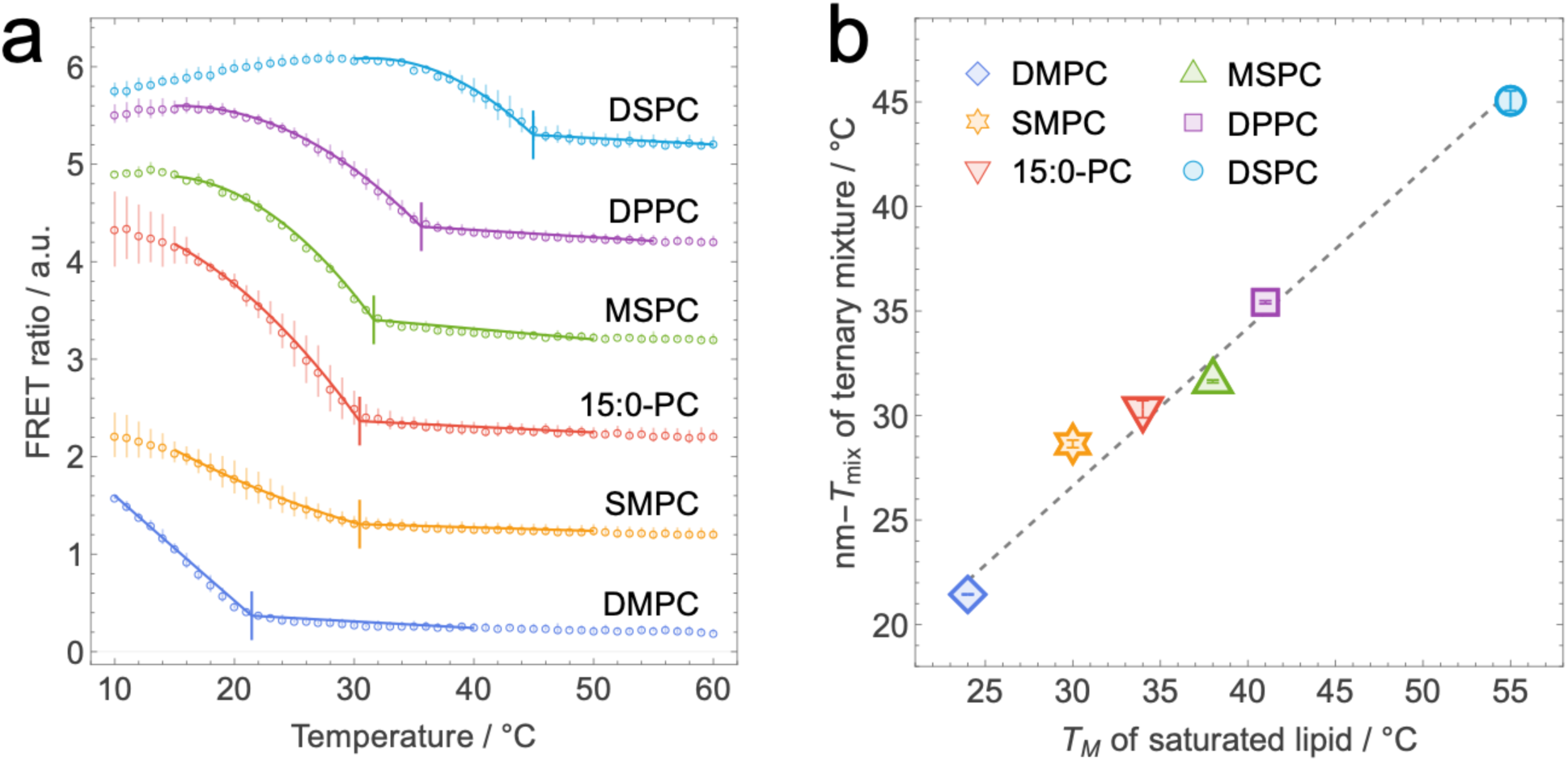
Nanoscopic miscibility transition temperature (nm-*T_mix_*) determined from FRET data. (a) Shown are FRET ratio vs. temperature data for ternary mixtures composed of high-*T_M_* lipid/DOPC/Chol 40/40/20 mol% for the indicated high-*T_M_* lipids. The solid lines are the best fit to a phenomenological model as described in Methods. Vertical lines indicate the best-fit value of nm-*T_mix_* determined from the model fits. Error bars are standard deviations determined from 10 sample replicates for SMPC/DOPC/Chol, 13 replicates for DPPC/DOPC/Chol, and 3 replicates for all other mixtures. (b) nm-*T_mix_* for ternary mixtures vs. the chain melting temperature, *T_M_*, of the saturated lipid. The dashed line is a linear fit (R^2^ = 0.999) to the data points corresponding to symmetric chain high-*T_M_* lipids (i.e., DMPC, 15:0-PC, DPPC, and DSPC). Error bars are SEM determined from sample replicates.

### 3.5. Comparing nm-T_mix_ and μm-T_mix_

Figure 7 plots nm-*T_mix_* vs. μm-*T_mix_* for the ternary mixtures, revealing a strong correlation between these transition temperatures. However, the two values are not identical; comparing the data points to the identity line (𝑦 = 𝑥, shown in light gray), it is clear that for each studied mixture, as the temperature is raised, sub-optical domains persist for another 4-11 °C after they are no longer visible by light microscopy (Table 2). This observation is in qualitative agreement with the 2D Ising model, which predicts the existence of nanoscopic compositional fluctuations above the microscopic miscibility critical point with a correlation length that decreases with increasing distance from the critical point [7, 50]. Our data are also consistent with a previously proposed mechanism in which a long-range repulsive interaction (for example, between lipid dipoles) competes with short-range attractive interactions to stabilize nanoscopic domains [51]. Domains with sub-optical dimensions are also predicted to arise from coupling between local composition and curvature [46, 47], however—as previously mentioned—it is difficult to reconcile the substantial bilayer thickness variation seen in cryo-EM images (Fig. 5) with the expected anti-registered domain arrangement predicted by the microemulsion theory. Still, we cannot rule out that the high curvature of extruded vesicles is somehow incompatible with a microemulsion. We stress that our data from compositionally symmetric model membranes does not allow us to comment on whether microemulsions are a plausible explanation for nanoscale heterogeneity in compositionally asymmetric membranes like the plasma membrane.

**Figure 7.**
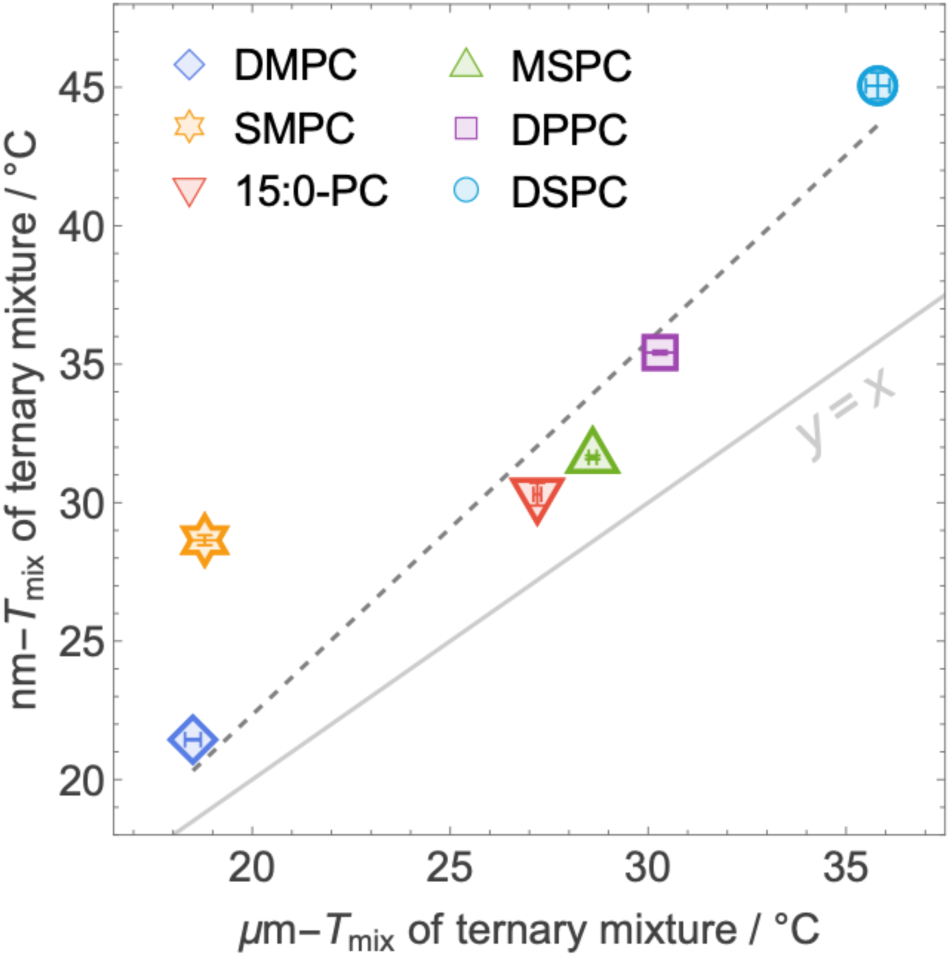
Comparing the microscopic and nanoscopic miscibility transition temperatures of ternary mixtures. Shown is the correlation between μm-*T_mix_* determined from fluorescence microscopy (horizontal axis) and nm-*T_mix_* determined from FRET (vertical axis) for ternary mixtures composed of high-*T_M_* lipid/DOPC/Chol 40/40/20 mol% for the indicated high-*T_M_* lipids. The dashed line is a linear fit (R^2^ = 0.998) to the data points for mixtures with symmetric chain high-*T_M_* lipids (i.e., DMPC, 15:0-PC, DPPC, and DSPC). The solid gray line corresponds to equality of nm-*T_mix_* and µm-*Tₘᵢₓ* values and is shown for reference.

The consistency of our finding that both macroscopic and nanoscopic domains exist in each of the studied systems, together with many similar observations both in model membranes [19, 52, 53] and GPMVs [54, 55] strongly suggests that sub-optical heterogeneity is universal in mixtures of low-and high-melting phospholipids plus cholesterol and does not depend on specific details of the system. Notably, none of the mixtures in our study contains a so-called hybrid lipid, i.e., one possessing both a saturated and an unsaturated chain. Clearly, such lipids are not a requirement for nanodomains [56], though they may help stabilize them by lowering the line tension at domain boundaries [57–59]. Along these lines, it is interesting that an unusually large gap between nm-*T_mix_* and μm-*T_mix_* occurs for the mixed-chain lipid SMPC. It is tempting to speculate that some mixed-chain saturated lipids might be capable of stabilizing the interface between ordered and disordered domains through a mechanism similar to that proposed for hybrid lipids. In this scenario, SMPC lipids that happen to be located at a boundary may show a preferred orientation, with the longer stearoyl chain residing primarily in the Lo domain and the shorter myristoyl chain residing in the Ld domain. The smaller gap between nm-*T_mix_* and μm-*T_mix_* observed for MSPC may be a consequence of the natural tilt of the glycerol backbone, which enhances the chain length difference for SMPC but diminishes it for MSPC. These hypotheses could be tested with molecular dynamics simulations.

Although our study focused on saturated mixed-chain PC lipids, a similar role could be played in eukaryotic PMs by sphingomyelins, some of which have a large length mismatch between their 16-carbon sphingosine chain and the N-acyl chain. Along these lines, Engberg and co-workers used FRET to measure nm-*T_mix_* of Lo domains in ternary mixtures containing low-melting lipids, cholesterol, and different sphingomyelins [60]. Interestingly, the largest and most thermostable domains were found in mixtures where the N-acyl chain closely matched the length of the sphingosine chain (i.e., 16 or 18 carbons). Reducing the length of the N-acyl chain to 14 carbons or increasing it to 24 carbons resulted in smaller and/or less abundant domains, suggesting a strong influence of chain length mismatch on domain properties in these mixtures.

## 4. Summary and Conclusions

We investigated ternary mixtures containing DOPC, cholesterol, and either symmetric-or mixed-chain saturated lipids. Our primary goal was to understand the influence of acyl chain length mismatch and backbone position within the high-melting lipid on the microscopic and nanoscopic phase behavior of these mixtures. Our findings can be summarized as follows:

1. For all studied mixtures, nm-*T_mix_* determined by FRET is 4-11 °C higher than μm-*T_mix_* determined by fluorescence microscopy, suggesting that nanoscopic heterogeneity is a universal feature that is not limited to mixtures containing hybrid low-*T_M_* lipids.
2. For all studied mixtures, there is a robust linear relationship between *T_M_* of the saturated lipid and nm-*T_mix_* of the mixture.
3. For *most* of the studied mixtures, μm-*T_mix_* is also strongly correlated with *T_M_*. The only exception is SMPC/DOPC/Chol, which has a μm-*T_mix_* that is substantially lower than expected from the trend exhibited by the other mixtures. This implies an expanded region of nanoscopic heterogeneity in SMPC/DOPC/Chol and suggests that the SMPC acyl chain structure is particularly conducive to forming nanoscopic domains, perhaps by preferentially aligning at domain boundaries and thereby lowering the interfacial energy. Interestingly, MSPC/DOPC/Chol adheres to the trend despite MPSC also having a four-carbon mismatch in chain length. We speculate that in this case, the natural tilt of the glycerol backbone blunts the effects of chain length mismatch.
4. Although fluorescence images of SMPC/DOPC/Chol GUVs are uniform at 22 °C, cryo-EM images of LUVs reveal coexisting thin and thick regions when the sample is cryo-preserved from 22 °C. This suggests that Ld and Lo domains are fully registered across the bilayer, in contradiction to an anti-registered alignment that would be expected for a 2D microemulsion originating from coupling of local composition and curvature.

## Author contributions

Deeksha Mehta: Conceptualization, Investigation, Formal analysis, Data curation, Visualization, Writing-Original draft, Writing-Review & editing

Emily Chaisson: Investigation, Formal analysis, Data curation, Visualization, Writing-Original draft, Writing-Review & editing

Averi M. Cooper: Investigation Maryam Ahmed: Investigation

M. Neal Waxham: Funding acquisition, Investigation, Data curation, Formal analysis, Writing-Review & editing

Frederick A. Heberle: Conceptualization, Funding acquisition, Supervision, Project administration, Formal analysis, Visualization, Writing-Original draft, Writing-Review & editing

## Supporting Information

The supporting information is available for the publication at (doi) and includes 4 figures.

## Acknowledgements

We are grateful to Dr. Tristan Bepler for providing an early version of MEMNET for automated contouring of vesicles in cryo-EM images, and to Mr. Venkata Mallampali for data acquisition assistance on the Titan Krios. Krios is partially supported by CPRIT Core Facility Grant RP190602. This work was supported by NIH grant R01GM138887 (to F.A.H. and M.N.W.). M.N.W. acknowledges the William Wheless III Professorship.

## References

[1] E. Sezgin, I. Levental, S. Mayor, C. Eggeling, The mystery of membrane organization: composition, regulation and roles of lipid rafts, Nat Rev Mol Cell Biol 18(6) (2017) 361–374.

[2] I. Levental, E. Lyman, Regulation of membrane protein structure and function by their lipid nano-environment, Nat Rev Mol Cell Biol 24(2) (2023) 107–122.

[3] K. Jacobson, P. Liu, B.C. Lagerholm, The Lateral Organization and Mobility of Plasma Membrane Components, Cell 177(4) (2019) 806–819.

[4] R. Kulkarni, E.A.C. Wiemer, W. Chang, Role of Lipid Rafts in Pathogen-Host Interaction - A Mini Review, Front Immunol 12 (2021) 815020.

[5] I. Levental, K.R. Levental, F.A. Heberle, Lipid Rafts: Controversies Resolved, Mysteries Remain, Trends Cell Biol 30(5) (2020) 341–353.

[6] F.A. Heberle, G.W. Feigenson, Phase separation in lipid membranes, Cold Spring Harb Perspect Biol 3(4) (2011).

[7] T.R. Shaw, S. Ghosh, S.L. Veatch, Critical Phenomena in Plasma Membrane Organization and Function, Annu Rev Phys Chem 72 (2021) 51–72.

[8] D. Marsh, Cholesterol-induced fluid membrane domains: a compendium of lipid-raft ternary phase diagrams, Biochim Biophys Acta 1788(10) (2009) 2114–23.

[9] G.W. Feigenson, Phase diagrams and lipid domains in multicomponent lipid bilayer mixtures, Biochim Biophys Acta 1788(1) (2009) 47–52.

[10] T. Harayama, H. Riezman, Understanding the diversity of membrane lipid composition, Nat Rev Mol Cell Biol 19(5) (2018) 281–296.

[11] T.M. Konyakhina, S.L. Goh, J. Amazon, F.A. Heberle, J. Wu, G.W. Feigenson, Control of a nanoscopic-to-macroscopic transition: modulated phases in four-component DSPC/DOPC/POPC/Chol giant unilamellar vesicles, Biophys J 101(2) (2011) L8–10.

[12] S.L. Veatch, S.L. Keller, Miscibility phase diagrams of giant vesicles containing sphingomyelin, Phys Rev Lett 94(14) (2005) 148101.

[13] P. Pathak, E. London, The Effect of Membrane Lipid Composition on the Formation of Lipid Ultrananodomains, Biophys J 109(8) (2015) 1630–8.

[14] T.A. Enoki, F.A. Heberle, G.W. Feigenson, FRET Detects the Size of Nanodomains for Coexisting Liquid-Disordered and Liquid-Ordered Phases, Biophys J 114(8) (2018) 1921–1935.

[15] I.V. Ionova, V.A. Livshits, D. Marsh, Phase diagram of ternary cholesterol/palmitoylsphingomyelin/palmitoyloleoyl-phosphatidylcholine mixtures: spin-label EPR study of lipid-raft formation, Biophys J 102(8) (2012) 1856–65.

[16] F.A. Heberle, J. Wu, S.L. Goh, R.S. Petruzielo, G.W. Feigenson, Comparison of three ternary lipid bilayer mixtures: FRET and ESR reveal nanodomains, Biophys J 99(10) (2010) 3309–18.

[17] F.A. Heberle, R.S. Petruzielo, J. Pan, P. Drazba, N. Kucerka, R.F. Standaert, G.W. Feigenson, J. Katsaras, Bilayer thickness mismatch controls domain size in model membranes, J Am Chem Soc 135(18) (2013) 6853–9.

[18] F.A. Heberle, M. Doktorova, H.L. Scott, A.D. Skinkle, M.N. Waxham, I. Levental, Direct label-free imaging of nanodomains in biomimetic and biological membranes by cryogenic electron microscopy, Proc Natl Acad Sci U S A 117(33) (2020) 19943–19952.

[19] D. Mehta, E.K. Crumley, J. Lou, B. Dzikovski, M.D. Best, M.N. Waxham, F.A. Heberle, Halogenated Cholesterol Alters the Phase Behavior of Ternary Lipid Membranes, J Phys Chem B 129(2) (2025) 671–683.

[20] J.H. Lorent, K.R. Levental, L. Ganesan, G. Rivera-Longsworth, E. Sezgin, M. Doktorova, E. Lyman, I. Levental, Plasma membranes are asymmetric in lipid unsaturation, packing and protein shape, Nat Chem Biol 16(6) (2020) 644–652.

[21] F.M. Goni, A. Alonso, Biophysics of sphingolipids I. Membrane properties of sphingosine, ceramides and other simple sphingolipids, Biochim Biophys Acta 1758(12) (2006) 1902–21.

[22] N. Jimenez-Rojo, A.B. Garcia-Arribas, J. Sot, A. Alonso, F.M. Goni, Lipid bilayers containing sphingomyelins and ceramides of varying N-acyl lengths: a glimpse into sphingolipid complexity, Biochim Biophys Acta 1838(1 Pt B) (2014) 456–64.

[23] M. Makarova, M. Peter, G. Balogh, A. Glatz, J.I. MacRae, N. Lopez Mora, P. Booth, E. Makeyev, L. Vigh, S. Oliferenko, Delineating the Rules for Structural Adaptation of Membrane-Associated Proteins to Evolutionary Changes in Membrane Lipidome, Curr Biol 30(3) (2020) 367–380 e8.

[24] D. Marsh, Structural and thermodynamic determinants of chain-melting transition temperatures for phospholipid and glycolipids membranes, Biochim Biophys Acta 1798(1) (2010) 40–51.

[25] M.P.K. Frewein, M. Doktorova, F.A. Heberle, H.L. Scott, E.F. Semeraro, L. Porcar, G. Pabst, Structure and Interdigitation of Chain-Asymmetric Phosphatidylcholines and Milk Sphingomyelin in the Fluid Phase, Symmetry (Basel) 13(8) (2021).

[26] E.G. Kelley, M.P.K. Frewein, O. Czakkel, M. Nagao, Nanoscale Bending Dynamics in Mixed-Chain Lipid Membranes, Symmetry (Basel) 15 (2023) 191.

[27] P.B. Kingsley, G.W. Feigenson, The synthesis of a perdeuterated phospholipid: 1, 2-dimyristoyl-sn-glycero-3-phosphocholine-d72, Chemistry and Physics of Lipids 24(2) (1979) 135–147.

[28] M.I. Angelova, D.S. Dimitrov, Liposome Electroformation, Faraday Discussions of the Chemical Society 81 (1986) 303–311.

[29] N.F. Morales-Penningston, J. Wu, E.R. Farkas, S.L. Goh, T.M. Konyakhina, J.Y. Zheng, W.W. Webb, G.W. Feigenson, GUV preparation and imaging: minimizing artifacts, Biochim Biophys Acta 1798(7) (2010) 1324–32.

[30] J.T. Buboltz, G.W. Feigenson, A novel strategy for the preparation of liposomes: rapid solvent exchange, Biochim Biophys Acta 1417(2) (1999) 232–45.

[31] E.H. Chaisson, D. Mehta, F.A. Heberle, Nanodomain formation in lipid bilayers I: Quantifying the nanoscopic miscibility transition with FRET, (2025).

[32] F.A. Heberle, M.N. Waxham, Phase separation in model lipid membranes investigated with cryogenic electron microscopy, Methods Enzymol 700 (2024) 189–216.

[33] F.A. Heberle, D. Welsch, H.L. Scott, M.N. Waxham, Optimization of cryo-electron microscopy for quantitative analysis of lipid bilayers, Biophys Rep (N Y) 3(1) (2023) 100090.

[34] A. Rohou, N. Grigorieff, CTFFIND4: Fast and accurate defocus estimation from electron micrographs, J Struct Biol 192(2) (2015) 216–21.

[35] R. Kiewisz, G. Fabig, W. Conway, J. Johnston, V.A. Kostyuchenko, A. Tan, C. Barinka, O. Clarke, M. Magaj, H. Yazdkhasti, F. Vallese, S.M. Lok, S. Redemann, T. Muller-Reichert, T. Bepler, Accurate and fast segmentation of filaments and membranes in micrographs and tomograms with TARDIS, bioRxiv (2025).

[36] S.C. Chen, J.M. Sturtevant, Thermotropic behavior of bilayers formed from mixed-chain phosphatidylcholines, Biochemistry 20(4) (1981) 713–8.

[37] T. Bultmann, H.N. Lin, Z.Q. Wang, C.H. Huang, Thermotropic and mixing behavior of mixed-chain phosphatidylcholines with molecular weights identical with that of L-alpha-dipalmitoylphosphatidylcholine, Biochemistry 30(29) (1991) 7194–202.

[38] S.W. Hui, J.T. Mason, C. Huang, Acyl chain interdigitation in saturated mixed-chain phosphatidylcholine bilayer dispersions, Biochemistry 23(23) (1984) 5570–7.

[39] C. Huang, J.T. Mason, Structure and properties of mixed-chain phospholipid assemblies, Biochim Biophys Acta 864(3-4) (1986) 423–70.

[40] G.J. Taylor, F.A. Heberle, J.S. Seinfeld, J. Katsaras, C.P. Collier, S.A. Sarles, Capacitive Detection of Low-Enthalpy, Higher-Order Phase Transitions in Synthetic and Natural Composition Lipid Membranes, Langmuir 33(38) (2017) 10016–10026.

[41] P.F. Almeida, Thermodynamics of lipid interactions in complex bilayers, Biochim Biophys Acta 1788(1) (2009) 72–85.

[42] R. Koynova, M. Caffrey, Phases and phase transitions of the phosphatidylcholines, Biochim Biophys Acta 1376(1) (1998) 91–145.

[43] C.E. Cornell, A. Mileant, N. Thakkar, K.K. Lee, S.L. Keller, Direct imaging of liquid domains in membranes by cryo-electron tomography, Proc Natl Acad Sci U S A 117(33) (2020) 19713–19719.

[44] K.D. Sharma, F.A. Heberle, M.N. Waxham, Visualizing lipid membrane structure with cryo-EM: past, present, and future, Emerg Top Life Sci 7(1) (2023) 55–65.

[45] P. Heftberger, B. Kollmitzer, A.A. Rieder, H. Amenitsch, G. Pabst, In situ determination of structure and fluctuations of coexisting fluid membrane domains, Biophys J 108(4) (2015) 854–862.

[46] S. Sadeghi, M. Muller, R.L. Vink, Raft formation in lipid bilayers coupled to curvature, Biophys J 107(7) (2014) 1591–600.

[47] D.W. Allender, M. Schick, A Theoretical Basis for Nanodomains, J Membr Biol 255(4-5) (2022) 451–460.

[48] M. Schick, Membrane heterogeneity: Manifestation of a curvature-induced microemulsion, Physical Review E—Statistical, Nonlinear, and Soft Matter Physics 85(3) (2012) 031902.

[49] M. Schick, Strongly Correlated Rafts in Both Leaves of an Asymmetric Bilayer, J Phys Chem B 122(13) (2018) 3251–3258.

[50] A.R. Honerkamp-Smith, S.L. Veatch, S.L. Keller, An introduction to critical points for biophysicists; observations of compositional heterogeneity in lipid membranes, Biochim Biophys Acta 1788(1) (2009) 53–63.

[51] R.D. Usery, T.A. Enoki, S.P. Wickramasinghe, M.D. Weiner, W.C. Tsai, M.B. Kim, S. Wang, T.L. Torng, D.G. Ackerman, F.A. Heberle, J. Katsaras, G.W. Feigenson, Line Tension Controls Liquid-Disordered + Liquid-Ordered Domain Size Transition in Lipid Bilayers, Biophys J 112(7) (2017) 1431–1443.

[52] S.L. Veatch, O. Soubias, S.L. Keller, K. Gawrisch, Critical fluctuations in domain-forming lipid mixtures, Proc Natl Acad Sci U S A 104(45) (2007) 17650–5.

[53] A.R. Honerkamp-Smith, P. Cicuta, M.D. Collins, S.L. Veatch, M. den Nijs, M. Schick, S.L. Keller, Line tensions, correlation lengths, and critical exponents in lipid membranes near critical points, Biophys J 95(1) (2008) 236–46.

[54] S.L. Veatch, P. Cicuta, P. Sengupta, A. Honerkamp-Smith, D. Holowka, B. Baird, Critical fluctuations in plasma membrane vesicles, ACS Chem Biol 3(5) (2008) 287–93.

[55] G. Li, Q. Wang, S. Kakuda, E. London, Nanodomains can persist at physiologic temperature in plasma membrane vesicles and be modulated by altering cell lipids, J Lipid Res 61(5) (2020) 758–766.

[56] F.A. Heberle, M. Doktorova, S.L. Goh, R.F. Standaert, J. Katsaras, G.W. Feigenson, Hybrid and nonhybrid lipids exert common effects on membrane raft size and morphology, J Am Chem Soc 135(40) (2013) 14932–5.

[57] T. Yamamoto, S.A. Safran, Line tension between domains in multicomponent membranes is sensitive to degree of unsaturation of hybrid lipids, Soft Matter 7(15) (2011) 7021–7033.

[58] Y. Hirose, S. Komura, D. Andelman, Concentration fluctuations and phase transitions in coupled modulated bilayers, Phys Rev E Stat Nonlin Soft Matter Phys 86(2 Pt 1) (2012) 021916.

[59] B. Palmieri, S.A. Safran, Hybrid lipids increase the probability of fluctuating nanodomains in mixed membranes, Langmuir 29(17) (2013) 5246–61.

[60] O. Engberg, K.L. Lin, V. Hautala, J.P. Slotte, T.K.M. Nyholm, Sphingomyelin Acyl Chains Influence the Formation of Sphingomyelin-and Cholesterol-Enriched Domains, Biophys J 119(5) (2020) 913–923.

