## Supporting Information for "Nanodomain formation in lipid bilayers II: The influence of mixed-chain saturated lipids"

[a]These authors contributed equally

**This PDF file includes:**

Figures S1-S4

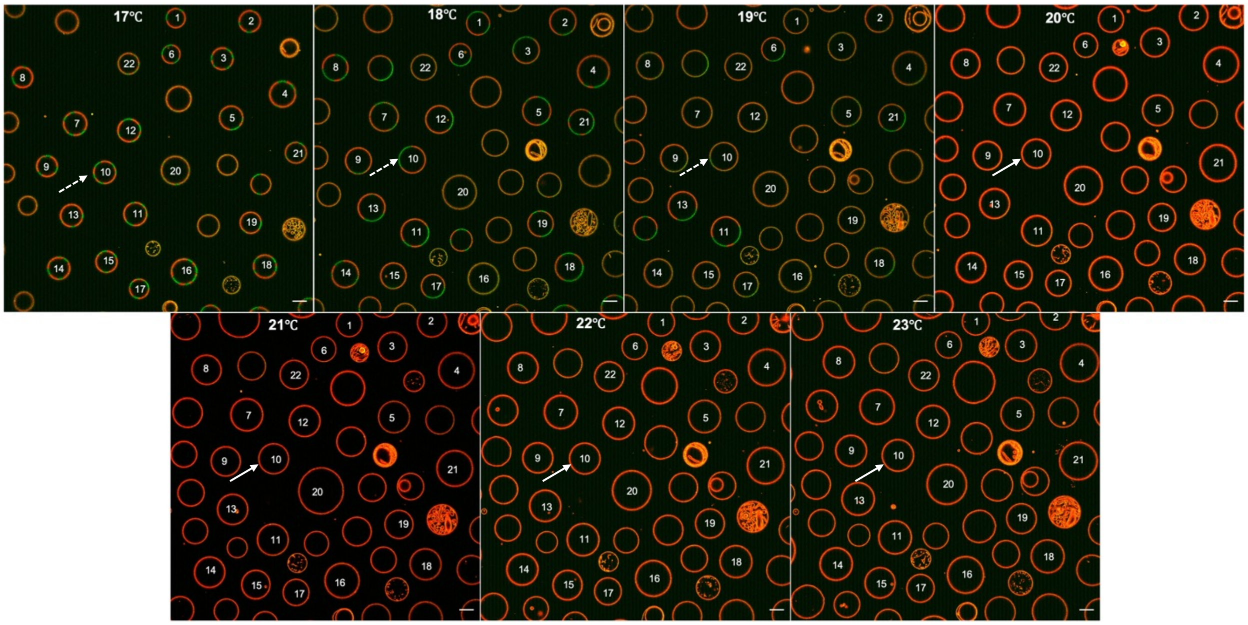


**Figure S1.** Determining the microscopic melting transition temperature from confocal fluorescence microscopy. Shown is a representative example of CFM images of GUVs used to determine µm-$T_{mix}$, here for SMPC/DOPC/Chol (40/40/20mol%). GUVs were labelled with fluorescent dyes LRPE (red) and NBD-DSPE (green), here shown as an overlay. The sample was initially cooled to a temperature where GUVs were phase separated (17 °C, upper left image) and then heated in 1 °C increments until all vesicles appeared uniform (23 °C, lower right image). The arrows follow one GUV through the miscibility transition, with dashed lines denoting the phase-separated state and solid lines denoting the uniform state. In this way, the fraction of phase separated GUVs is determined each temperature and then fit to a sigmoidal function to determine µm-$T_{mix}$. Scale bars are 10 µm.

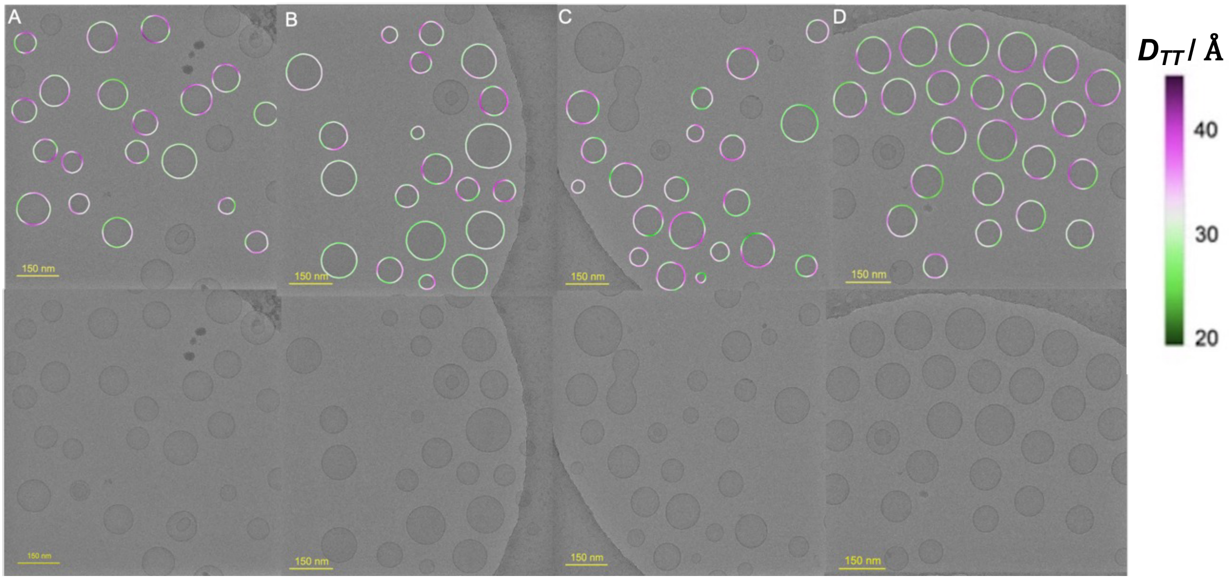


**Figure S2.** Full field of view cryo-EM images of ternary liposomes. Top and bottom rows are the same image without (bottom) and with (top) a color overlay corresponding to the bilayer thickness, $D_{TT}$. Vesicle compositions (40/40/20 mol%) are: DPPC/DOPC/Chol (A); (B) MSPC/DOPC/Chol (B); 15:0-PC/DOPC/Chol (C); and SMPC/DOPC/Chol (D). Vesicles highlighted with color overlays in the top row were selected for analysis as described in Methods. Scale bars are 150 nm.

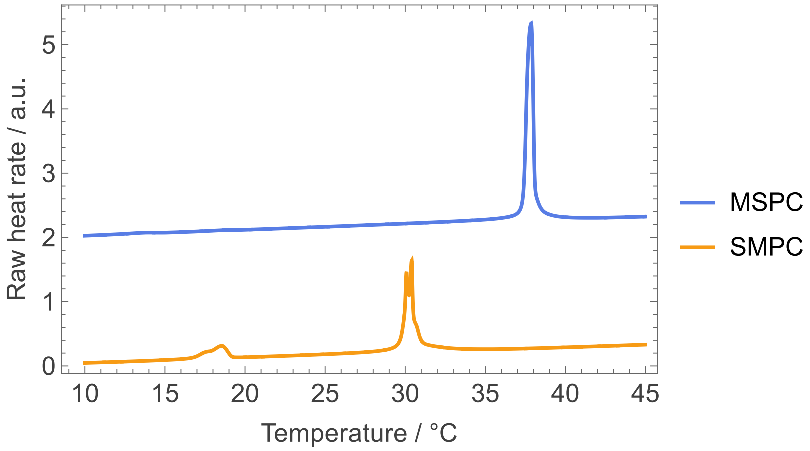


**Figure S3.** Differential scanning calorimetry (DSC) thermograms (heating scans) of SMPC and MSPC. The measured transition temperatures and peak shapes closely match those reported by Chen and Sturtevant for these lipids (ref. 36 in the main text), indicating a comparable extent of positional isomerization and confirming the expected thermotropic behavior of the mixed-chain PCs.

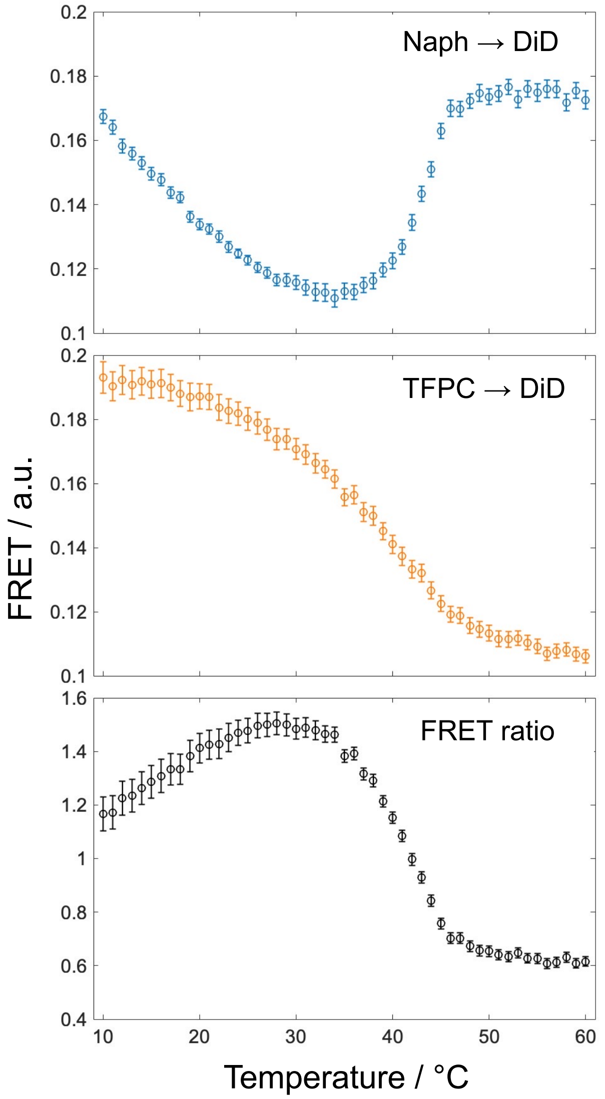


**Figure S4.** FRET data for 40/40/20 mol% DSPC/DOPC/Chol. Shown is the normalized sensitized acceptor emission for Nap donor to DiD acceptor (top), TFPC donor to DiD acceptor (middle), and the ratio of these signals (bottom).
